# Planktonic oyster larvae optimize settlement decisions in complex sensory landscapes

**DOI:** 10.1101/2024.11.28.625415

**Authors:** Sarah Schmidlin, Yliam Treherne, Jan Mees, Pascal I. Hablützel

## Abstract

The settlement of pelagic larvae constitutes a pivotal phase in the life cycle of benthic aquatic species. The choice of settlement location is critical for the recruitment to established populations and the colonization of unoccupied habitats. Consequently, the cues governing settlement decisions in larvae become particularly pertinent for human activities such as habitat restoration, mariculture, or biofouling prevention. Our study aims to enhance our comprehension of the underlying principles of how larvae optimize settlement decisions when exposed to multiple natural chemical cues simultaneously. Through a series of laboratory experiments, we investigate settlement patterns of Pacific oyster *Magallana gigas* larvae when exposed to different combinations of attractant and repellent cues. Our findings reveal additive increases in settlement rates in the presence of attractant cues originating from conspecifics and biofilms. Conversely, settlement attraction by conspecific water-borne cues was reduced in the presence of repellent cues emanating from predators. Notably, when repellent predator cues were presented alongside attractant cues linked to substrates (biofilms or shells from conspecific adults), the repellent effect was nullified.

## Introduction

Settlement is a critical step in the life history of aquatic species with a planktonic larval stage. In sessile organisms, the process of settlement consists of descent from the water column to the substrate surface, attachment, and finally metamorphosis (Morse., 1990; Hadfield & Paul 2001). For organisms with only limited or no relocation ability this process is essential for ultimate survival of a population (Crisp 1976, Bayne 2017). Settling larvae, therefore, screen their environment for cues that will give away information about the suitability of a site for settlement and survival (Zimme-Faust & Tamburri 1994, Hadfield & Paul 2001). Such information may include abiotic conditions (e.g. salinity, light, water current; Rittschof et al., 1998; Koehl, 2007; Kim et al., 2021), presence of conspecifics (Hadfield & Paul 2001, Dobretsov & Rittschof 2020), predators (Pruett & Weissburg 2019, Scrosati 2021), and biofilms (Hadfield 2011, Campbell et al. 2011, Dobretsov & Rittschof 2020), or the availability of food (Forêt et al. 2018). Depending on the information they carry, cues can elicit different reactions by being either attractive or repellent. Specific environments may release different cues at once, sending potentially conflicting messages to the pelagic larvae. For settling larvae, the optimal behavioral strategy would be to respond not only to cues that signal immediate risks and opportunities but also to evaluate future conditions for feeding or mating. To maximize survival and fitness, the larvae should therefore assess the relative importance of multiple and potentially conflicting cue for current and future survival, growth, and reproduction and optimize (optimality theory (Milinski & Heller 1978, Sih 1980)) their settlement decision depending on the relative importance of the cues (threat sensitivity hypothesis (Sih 1986, Helfman 1989)).

In this study, we seek to advance our understanding of how multiple natural chemical cues interact to influence settlement behaviors for pediveliger larvae of *M. gigas*. Specifically, how larvae behave when facing simultaneous signaling from contradicting attracting (conspecific adults) and repelling (predators) cues. When faced with multiple conflicting cues i.e. those associated with conspecifics and those associated with predators, we propose that larvae could adopt the following decision rules: (1) if the positive cue is perceived as important, larvae settle when exposed to positive cues regardless of the presence of a negative cue, (2) if the negative cue is perceived as important, larvae delay settlement in the presence of negative cues regardless of the addition of the positive cue, (3) in the absence of a specific preference, larvae add up positive and negative cues and the larvae’s response depends on their relative strength. To test these hypotheses, we exposed lab-reared pediveliger larvae to conspecific, predator and biofilm cues in full-factorial no-choice experiments. Conspecific and biofilm cues were included as they are well known positive inducers of oyster settlement (Tamburri, Zimmer-Faust, & Tamplin., 1992). Predator cues have previously been shown to decrease oyster settlement (Pruett & Weissburg 2019). In response to predator kairomones, larvae of invertebrates may display avoidance or protection behaviors such as orienting swimming direction away from predator direction (Morello & Yund, 2016), increasing growth rates (Pruett & Weissburg 2019), or reduced settlement propensity (Bertolini et al., 2019).

## Methods

### Overview

We conducted two laboratory experiments to assess the effect of various putative settlement cues and their interaction on settlement propensity in Pacific oyster larvae. Experiments took place in Petri dishes where competent pediveliger larvae were exposed for 30 hours to different combinations of positive and negative settlement cues in full factorial designs. In the first experiment (experiment 1), performed in May 2022, we subjected larvae to three different cues in a full-factorial design. These cues were conspecific waterborne cues, conspecific shell cues, and waterborne cues from natural predator *Carcinus maenas*. We replicated all treatments in 20 separate Petri dishes, with 6 larvae per dish, over 2 consecutive days with 10 replicates performed each day.

Based on the results of experiment 1, we performed a second experiment (experiment 2) in August 2022 using cues from predator *C. maenas*, from biofilms, and from both waterborne and shell-associated conspecific cues. In experiment 2, we considered 16 different treatment combinations in a fully factorial design. We repeated each treatment combination in 25 separate Petri dishes, with 6 larvae per dish, over 3 consecutive days with 8 replicates of each treatment combination being performed each day. A smaller pilot study was performed in March 2022. This pilot study was not included in the final analysis as cues were not added in similar volumes as the subsequent experiments. All information related to the pilot study can be found in the supporting information table and figure S1.

### Larvae cultures

The parent oysters used in experiment 1, were purchased from the Guernsey Sea Farms Ltd (Guernsey, UK). We fertilized two batches of eggs from these adults (five of each sex) five days apart so that competent larvae were available during all days of the experiment. For experiment 2, we used mature adult oysters (10 of each sex) from the harbor of Ostend (Belgium). The process of rearing larvae was kept the same in both experiments. See supplement for a detailed account. Immediately prior to using larvae in the experiments, larvae were taken from the culture tanks and filtered through two consecutive sieves of 300 and 260 μm. Only larvae retained on the second (260 μm) sieve were used in the experiment. Larvae were allowed to acclimate in filtered sea water (FSW) for an hour in the climate-controlled room where experiments took place.

### Preparation of waterborne cues

Production of all waterborne cues (predator and conspecific) was performed similarly to the methods from Morello and Yund (2016). For production of conspecific cues, adult oysters were collected (wet weight of soft parts was 7.6 ± 2.96 g) from the harbor of Ostend (Belgium) 3 days prior to the start of the experiments. Adults were scrubbed clean of any biofilms and kept for 24 hours in filtered seawater at 15 °C. Subsequently, we placed eight adult oysters in 1800 ml of filtered aerated water for 48 hours and then filtered the water through a 30 µm nylon sieve. Water was kept at 15 °C until used in experiments.

### Preparation of Predator cues

To produce predator cues, we collected adult individuals of *C. maenas* from the harbor of Ostend (Belgium) and held them in FSW aerated tanks at 15 °C. Predator cue water was created similarly to conspecific cue by placing one crab into 600 ml of FSW for 48 hours. Four crabs were used in the first experiment and three crabs were used in the second. The average weight of crabs in the first experiment was 53.2 ± 13 g and measuring 6.05 ± 0.62 cm in max. carapace length in the first experiment. In the second experiment crabs were 66.8 ± 5g and 6.6 ± 0.60 cm were used and cue water was combined before use in experiments. At 48 hours crabs were removed and cue water was filtered through a 30 μm sieve.

### Preparation of conspecific shell cues

Conspecific shell chips came from live oysters collected from the Ostend coast. First, we removed the soft parts of the oyster and scrubbed the shells thoroughly. We then crushed the shells using a hammer and sieved them through 1.0 mm and 0.5 mm metal sieves, collecting the shell fragments that were retained by the 0.5 mm mesh screen and drying them in a 30 °C oven. Shells were stored in a freezer at −20 °C until use in experiments (similar to the method used by Vasquez et al., (2013)). To create substrates without an active cue, we heated the shell chips at 300 °C for 3 hours. This has been shown in previous research to eliminate the attractive properties of the cue associated with the shell (Vasquez et al., 2013). The preparation of chips without attraction was necessary so that all treatments would have the same settlement surface topography.

### Preparation of biofilm cues

Natural biofilms were allowed to develop on shell chips placed near the surface in the non-tidal part of the Ostend harbor. Shell chips were kept in 300 μm nylon mesh, allowing water to flow through to the shell chips, enabling colonization by microbial organisms. Biofilms were left to develop undisturbed for 9 days. Immediately prior to experiments we retrieved the chips with biofilms and transported them with natural seawater to prevent them from drying out.

### Settlement Assessment

Per replicate, we moved six random pediveliger larvae into 15 ml Petri dishes with shell chip substrate (either substrate with cue or sham chips without cue). Each Petri dish contained 7 ml of one or both of the two types of treatment water and were topped up with FSW for a total of 14 ml. Petri dishes were made of polystyrene and had a diameter of 35 mm and a maximum volume of 15 ml. New plates were used for each replication. We evaluated the settlement cue treatments by counting the number of individuals that metamorphosed to settled post-larvae after 30 hours of exposure. In experiment one settlement was assessed after 10, 20, and 30 hours, larvae settlement increased in all treatments overtime but and no differences were noted in interactions. Due to the low settlement at shorter time allowed to settle, statistical analysis was more difficult, thus 30 hours was chosen as the time to assess settlement (see table S2 in supplement). Microalgae were added to the seawater in each treatment at the same rate as larvae cultures (*Caetoceros muelleri*, and *Isochrysis galbana* at 100,000 cells/ml at a volume ratio of 3:1) and were not limiting throughout the duration of the experiment. All trials happened in a 12 h day, 12 h night environment at 19 (±1) °C in a climate-controlled room.

### Statistical analyses

For each experiment we created a set of generalized linearized mixed-effect models using the glmer function of the lme4 package (Bates et al., 2014) in R version 4.1.3 (2022-03-10) (R Core Team, 2021). As the response variable was binary (settled vs. not settled) we fitted a Bernoulli distribution using a logit link function. For each experiment, a base model was established, consisting of each cue treatment (conspecific cue, predator cue, conspecific shell, and biofilm) as fixed effect variables, and the batch from which the larvae originated as a random variable. A forward selection procedure, using the Akaike Information Criterion (AIC), was performed to determine if 1) any interaction effects of cue treatments or 2) the age of the larvae should also be included as fixed effect variables in the model. A description of the final models can be found in the supporting information. As experiments 1 and 2 had the same methodology and treatments, we combined the data in a final analysis to increase statistical power. Finally, we performed post-hoc tests using the emmeans function in R to calculate the marginal means adjusting p-values for multiple comparisons with Tukey’s method and used the pairs function to display pairwise comparisons. See supporting information for model descriptions and descriptions of post-hoc comparisons (tables S3 and S4). Results display prediction plots produced by the model using the ggpredict package in R. The effect of the treatments is given in predicted percent settled from the model.

## Results

### Experiment 1

In the first experiment, Predator cues did not decrease settlement significantly, however there was a non-significant trend showing a reduction of settlement (p=0.09) (Table 1, Figure 1). The presence of conspecific waterborne cues and conspecific shell both significantly increased predicted settlement (Table 1, Figure 1). Post-hoc tests showed that positive cues from conspecific cues significantly increased settlement in the waterborne cue and shell cue combined treatment (see also Table S4 in supplement), but as there was no interaction effect, and we cannot say that this was a synergistic interaction between positive cues (Table 1).

**Table 1.** Results of the statistical models for each experiment. Significant (p < 0.05) are in bold, while marginally-significant (p < 0.1) are in *italic*.

| Experiment | Predictor Variable | Estimate | Std. error | z-value | p-value |
| --- | --- | --- | --- | --- | --- |
| 1 | Conspecific cue present | 1.59795 | 0.52913 | 3.020 | **0.00253** |
|  | Predator cue present | -1.86435 | 1.10798 | -1.683 | *0.09244* |
|  | Shell cue present | 3.28470 | 0.51480 | 6.380 | **1.77e-10** |
|  | Interaction<br>conspecific cue -<br>predator cue | 0.94550 | 1.18659 | 0.797 | 0.42555 |
|  | Interaction<br>conspecific cue -<br>shell cue | -0.23820 | 0.63500 | -0.375 | 0.70757 |
|  | Interaction<br>predator cue -<br>shell cue | 2.18025 | 1.15149 | 1.893 | *0.05830* |
| 2 | Conspecific cue<br>present | 1.78016 | 0.19217 | 9.263 | **< 2e-16** |
|  | Predator cue<br>present | 0.20793 | 0.18838 | 1.104 | 0.2697 |
|  | Shell cue present | 2.24582 | 0.16348 | 13.738 | **< 2e-16** |
|  | Biofilm present | 1.31959 | 0.23868 | 5.529 | **3.22e-08** |
|  | Interaction<br>conspecific cue -<br>predator cue | -0.42616 | 0.21340 | -1.997 | **0.0458** |
|  | Interaction<br>conspecific cue -<br>biofilm | -0.45345 | 0.22080 | -2.054 | **0.0400** |
|  | Interaction shell<br>cue present -<br>biofilm | -0.40334 | 0.22253 | -1.812 | *0.0699* |
|  | Interaction<br>predator cue - biofilm | -0.26063 | 0.21334 | -1.222 | 0.2218 |
| combined | Conspecific cue present | 1.51893 | 0.13632 | 11.142 | **< 2e-16** |
|  | Predator cue present | 0.06348 | 0.13248 | 0.479 | 0.63181 |
|  | Shell cue present | 2.33891 | 0.10274 | 22.765 | **< 2e-16** |
|  | Interaction conspecific cue - predator cue | -0.55469 | 0.25121 | -2.208 | **0.0272** |
|  | Interaction conspecific cue – Shell | -0.49814 | 0.19050 | -2.615 | 0.00892 |

**Figure 1.**
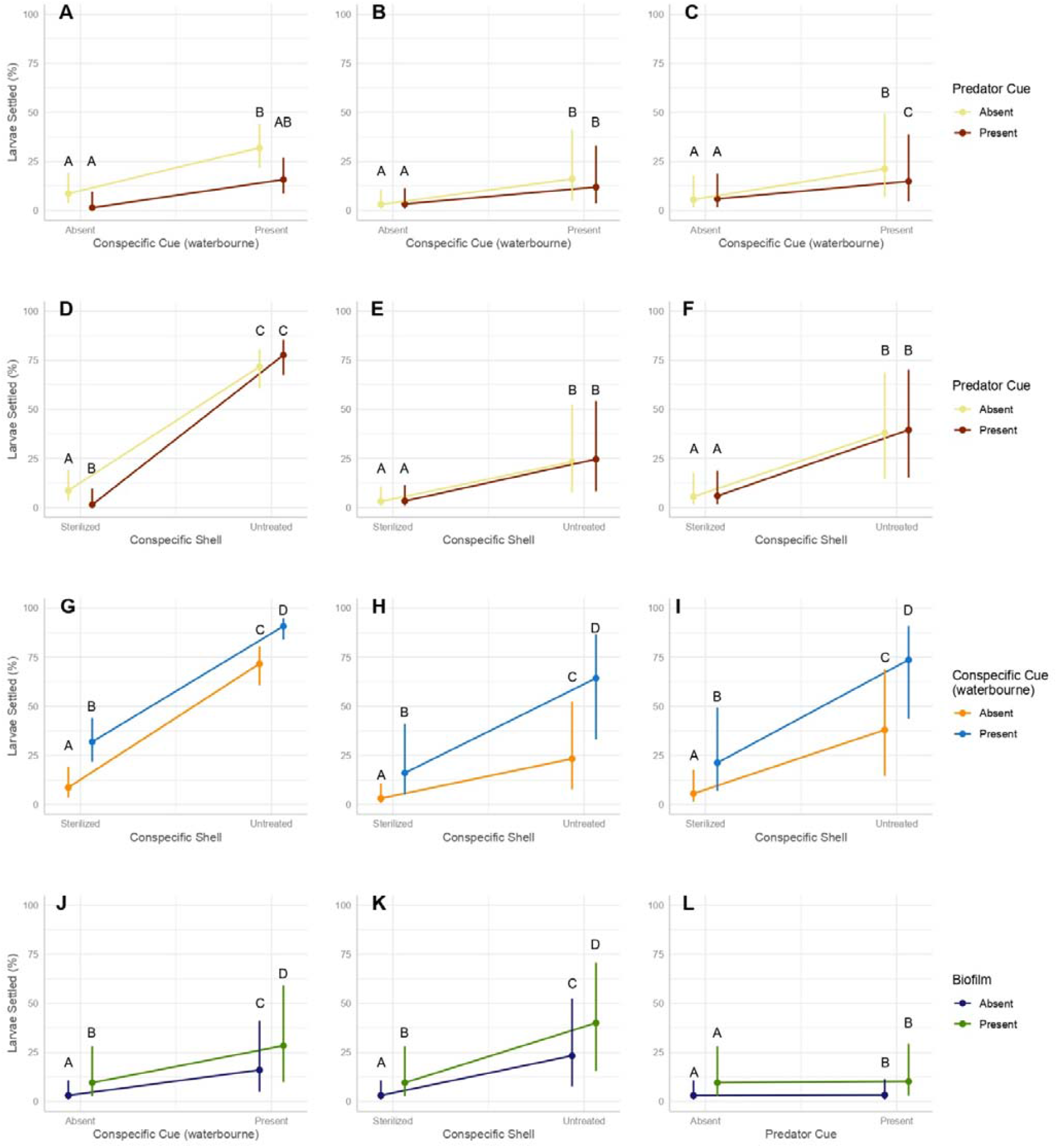
Predictions of the generalized linear mixed effect model showing estimated probability of larvae settlement. Error bars represent 95 % confidence intervals of model prediction. Images A, B and C show predicted larvae settlement when exposed to cues from predator *C. maenas* and conspecific waterborne cues. Image A shows results from experiment 1, image B shows results from experiment 2 and image C shows results from combined experiment data. Images D, F, and E show predicted larvae settlement when exposed to cues from predator *C. maenas* and conspecific shells. Image D shows results from experiment 1, image E shows results from experiment 2 and image F shows results from combined experiment data. Images G, H, and I show predicted larvae settlement when exposed to cues from waterborne and shell conspecific cues image G shows results from experiment 1, image H shows results from experiment 2 and image I shows results from combined experiment data. Image J shows predicted settlement when exposed to cues from waterborne conspecific cues and biofilms, image K shows predicted settlement from conspecific shells and biofilms, and image L shows predicted settlement from predator cues and biofilms. Results from J, K, and L come from experiment 2.

### Experiment 2

In the second experiment, the presence of waterborne conspecific cues, conspecific shells, and biofilms significantly increased the predicted settlement (Table 1). Post-hoc tests showed that for all positive cues, settlement is significantly increased with each additional cue (Figure 1). Significant interaction between waterbourn conspecific cues and biofilm cues indicate some synergistic relationship (Table 1). The presence of predator cues did not significantly change the predicted settlement (Table 1). Predator cues did not decrease settlement significantly when exposed to the presence of conspecific waterborne cues, biofilms, and conspecific shell but in the case of waterborne conspecific cues and predator cues together there was a non-significant trend for some reduction in settlement (Figure 1) (see also table S4 in supplement).

### Combined

In the combined analysis results from conspecific cues remained consistent (Table 1, Figure 1). Significant interaction between waterbourn conspecific cues and shell cues indicate some synergistic relationship (Table 1). Post-hoc tests reveal that when conspecific waterborne cues and predator cues are both present, the presence of predator cues decreases settlement significantly (Figure 1, odds ratio = 1.544, P = 0.0081) (see post-hoc description in supplement table S4). When positive cues from conspecific shells were present together with predator cues, settlement was not reduced (Figure 1, odds ratio = 1.204, P = 0.207) (see also table S4 in supplement).

## Discussion

Natural environments are complex sensory landscapes. Multiple simultaneous settlement cues may require the larva’s responses to an individual cue to change depending on the prevailing combination of cues. However, little is known about behavioral responses to multiple conflicting cues. In the present study, we found that in response to conflicting cues from a combination of waterborne positive conspecific cues and negative predator cues oyster larvae will decrease settlement, compared to a treatment with only conspecific cues. Interestingly, we did not observe a reaction to predator kairomones in the presence of positive cues from oyster shells or biofilms. While previous work has identified (or theorized) that environmental cues may be ranked in a hierarchical way (Igulu et al., 2013; Hodin et al., 2018) to the best of our knowledge these are the first results showing that this hierarchy applies to predator cue interpretation. Discrimination of cues may play an important role in decision making in the presence of multiple conflicting cues.

### Settling larvae may rank conflicting cues

Our results indicate that when exposed to conflicting cues, *M. gigas* larvae adjust their behavior in a way that is specific to the respective combination of cues. When presented with two conflicting waterborne cues from conspecifics and their predator *Carcinus maenas*, it seems that larvae added up positive and negative cues and weighed their decision for settlement accordingly. This means that the presence of predator kairomones reduced settlement, but not to the level observed in the absence of the positive cue. But when faced with positive cues associated with the shells of conspecifics or with biofilms, larvae were less risk averse and settled regardless of the presence of a negative cue. This observation could suggest that oyster larvae may perceive cues differently based on their source (waterborne vs substrate-bound). If the positive cue is perceived as stronger or more important than the negative cue, the negative cue will be disregarded (or at least downweighed), resulting in an optimistic decision for settlement. The cues we found in this study to be high ranking were those associated with conspecific shells and biofilms, both of which are substrate-bound. In the case of a similar ranking, the additive effect of both cues will render the settlement choice balanced according to the relative strength of the cues, as seen in the conflicting waterborne cues in our experiment. Extrapolating, this would imply that for the lowest ranks of positive cues, the larvae use a pessimistic strategy and do not settle in the presence of negative cues. Combinations of multiple positive cues were seen to increase settlement in an additive way: the addition of simultaneous positive cues will provide more confidence of larvae in their settlement choice. While adding multiple cues increases confidence, the observed relationship between these cues in this study was additive, meaning that multiple cues increase the chance of settlement, but not synergistic, meaning that the presence of a second positive cue does not increase the effect of the first.

### Settlement cues in a small-scale experimental setup

In our study, we used 15 ml Petri dishes for settlement trials. In such small volumes contact with the substrate is unavoidable. Therefore, we cannot discriminate between the effect of settlement cues on initiating descent from the water column to the substrate and on metamorphosis, respectively. In many marine invertebrate taxa, both the initial larva descent to a substrate, also sometimes referred to as settlement, and metamorphosis can be induced by a single cue source (Morse and Morse, 1984; McGee and Targett, 1989; Pearce and Scheibling, 1990). Larvae of other taxa require a combination of waterborne and substrate-bound cues to first “settle” and then metamorphize (Chia and Koss, 1988). Different signaling pathways may exist in bivalve larvae for perceiving and reacting to cues (Wang et al., 2023). The scope of acceptable cues for bivalve settlement and metamorphosis tends to be broad, and which combination is needed can be species specific (Bao et al., 2007; Yang et al., 2013). For M. gigas, waterborne conspecific cues have been shown to increase settlement propensity (Zimme-Faust & Tamburri, 1994) and – as indicated in our study – also metamorphosis. One could argue that waterborne cues become irrelevant once larvae sense a substrate-bound cue. Presenting larvae with waterborne and substrate-bound cues in a 15 ml Petri dish may not be ecologically realistic. While keeping this limitation in mind, we argue that waterborne cues may still be relevant in our setup as the larvae’s propensity to settle further increased when provided with positive waterborne cues in addition to positive substrate-bound cues. While waterborne cues can be distributed in the water column beyond the limits of the reef, substrate-bound cues indicate an immediate favorable settlement location. As such, the waterborne cues may play a more important role in inducing initial settlement, while at a later stage, larvae probe the substrate for further substrate-bound cues that induce metamorphosis. Our results would indicate that after contact with a substrate-bound cue, predator cues lose importance for the decision to metamorphize.

### Implications for oyster reef management

Settlement is a critically important life history phase because this transition is irreversible and because wrong choices can reduce survival, growth and reproductive success (Jenkins et al., 2009). Marine invertebrate larvae have evolved to make the transition from pelagic to sessile quickly once suitable conditions are met, in contrast to other metamorphosing taxa like insects and amphibians (Hadfield, 2000). This may be especially true for Pacific oyster *M. gigas*, which has broader ecological tolerances than other oyster species (Troost, 2010). For example, oyster *Ostrea edulis* reproduces through internal fertilization (Bayne, 2017), meaning that proximity to conspecific adults may be perceived as more critical than for *M. gigas* which are external broadcast spawners. For generalist species like *M. gigas*, it therefore might not be very advantageous to delay settlement until suitable conditions are met, thus favoring the evolution of optimistic settlement strategies. In the long run, insights from this research may enable ecosystem management practitioners when designing reef restoration projects, spat collectors for bivalve mariculture, or developing biofouling mitigation strategies.

## Supporting information

Supplement to paper_Planktonic oyster larvae optimize settlement decisions in complex sensory landscapes

## Acknowledgments

We thank the Research Foundation - Flanders (FWO) in the framework of the Flemish contribution to LifeWatch, which is a landmark European Research Infrastructures on the European Strategy Forum on Research (ESFRI) roadmap for providing infrastructure. We would also like to thank Mattias Bossaer for technical assistance and Dr. Nancy Nevejan for advice on oyster larviculture. This article was improved by the valuable comments of two anonymous reviewers.

