## Supplement to paper_Planktonic oyster larvae optimize settlement decisions in complex sensory landscapes for "Planktonic oyster larvae optimize settlement decisions in complex sensory landscapes"

This is the supporting information for the manuscript titled “Planktonic oyster larvae optimize settlement decisions in complex sensory landscapes”

**Pilot study**

In a pilot study, performed in March 2022, larvae were subjected to four different treatments of cues and cue combinations; conspecific waterborne cue only, predator waterborne cue from *C. maenas* only, conspecific waterborne cue + predator (*C. maenas)* waterborne cue, and a control treatment without cues. This pilot study was not included in the final analysis as cues were not added in similar volumes in this study. Waterborne conspecific cues and predator cues were added at 14ml to each petri dish in isolated treatments but in treatments where cues were combined there were 7ml of conspecific waterborne cues and 7ml of predator cue. Preparation of the waterborne conspecific cue and the predator cues were the same as those listed in the main experiments.

Statistical analyses were performed similar to the main experiments, a generalized linearized mixed-effect model was used to compare the interaction of conspecific waterborne cue and predator cue (Bates et al., 2015) in R version 4.1.3 (2022-03-10) (R Core Team, 2021).

Table S1. Results of the statistical model for the pilot experiment

| **Predictor variable** | **Estimate** | **Std. error** | **z-value** | **p-value** |
| --- | --- | --- | --- | --- |
| Conspecific cue present | 1.5411 | 0.39488 | 3.903 | **9.51E-05** |
| Predator cue present | 0.48187 | 0.44092 | 1.093 | 0.27445 |
| Interaction conspecific cue - predator cue | -1.28398 | 0.54041 | -2.376 | **0.01750** |

Table S2. Experiment 1 results from metamorphosis assessed after 10, 20, and 30 hours.

| Hour assessed | **Predictor Variable** | **Estimate** | **Std. error** | **z-value** | **p-value** |
| --- | --- | --- | --- | --- | --- |
| 30 hrs | Conspecific cue present | 1.59795 | 0.52913 | 3.020 | **0.00253** |
|  | Predator cue present | -1.86435 | 1.10798 | -1.683 | *0.09244* |
|  | Shell cue present | 3.28470 | 0.51480 | 6.380 | **1.77e-10** |
|  | Interaction conspecific cue - predator cue | 0.94550 | 1.18659 | 0.797 | 0.42555 |
| 20hrs | Conspecific cue present | 2.27899 | 0.76780 | 2.968 | **0.003** |
|  | Predator cue present | -0.90030 | 1.23592 | -0.728 | 0.466 |
|  | Shell cue present | 3.59219 | 0.74970 | 4.792 | **1.66e-06** |
|  | Interaction conspecific cue - predator cue | 0.33131 | 1.30703 | 0.253 | 0.800 |
| 10hrs | Conspecific cue present | 1.75764 | 0.78663 | 2.234 | **0.025458** |
|  | Predator cue present | -14.80977 | 124.18193 | -0.119 | 0.905070 |
|  | Shell cue present | 2.54814 | 0.75966 | 3.354 | **0.000796** |
|  | Interaction conspecific cue - predator cue | 13.88916 | 124.17916 | 0.112 | 0.910944 |

**Figure S1**

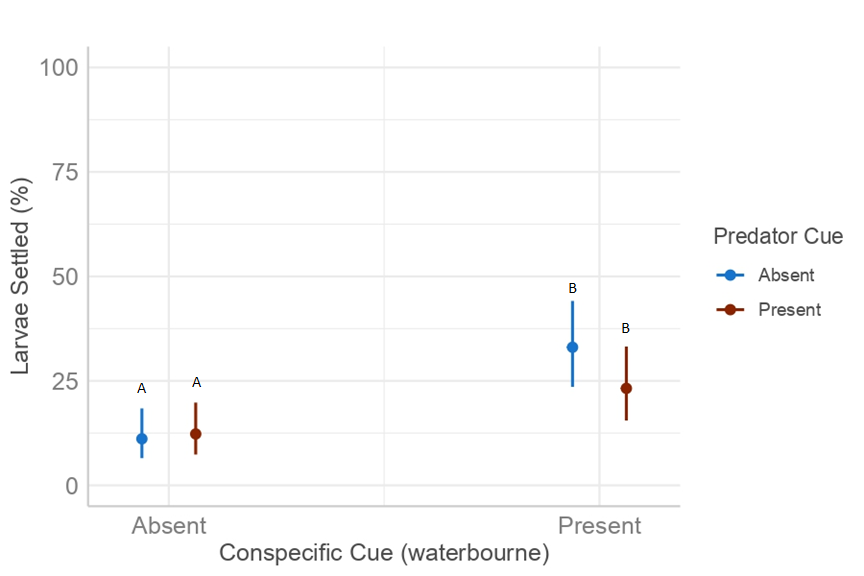

Figure S1. Predictions of the generalized linear mixed effect model showing estimated probability of larvae settlement when exposed to predator cues from *C. maenas* and conspecific waterborne cues. Error bars represent 95 % confidence intervals of model prediction. This graph represents the results of the Pilot study showing interaction of waterborne conspecific cues and predator cues from *C. maenas.*

**Larval Rearing**

Adult oyster were induced to spawn through gonad stripping, and eggs were fertilized following FAO guidelines (Helm, 2004). Fertilized eggs were kept undisturbed in 5-liter flat bottom tanks for 48 hours at 22 °C at a density of 10 eggs per ml of 0.1 μm filtered seawater (FSW). All seawater was sourced from the water basin at NIOZ Yerseke and filtered at 0.1 μm to remove harmful bacteria and other small microorganisms. After 48 hours larvae were sieved over 30 µm nylon mesh, rinsed, and added larvae to 10-liter conical tanks with FSW. Conical tanks were static but aerated and kept at 22 °C for the entire duration of larvae rearing. Every two days, we sieved the larvae over mesh corresponding to the average size of the larvae and the water in the tanks was changed. Seawater used throughout the larval rearing and experiment maintained a pH of 7.8 ± 0.95 and salinity of 31.4 ± 1.2 in the first experiment and pH of 8.03 ± 0.05, and salinity of 32.7 ± 0.49 in the second experiment. Larvae fed *ad libitum* from a fresh microalgae mixture consisting of *Chaetoceros muelleri*, and *Isochrysis galbana* (clone T-ISO) purchased from Proviron Industries NV. For the first four days larvae were fed at 40,000 cells/ml water using only *I. galbana*. Days 5-12 larvae were fed *C. muelleri*, and *I. galbana* at 100,000 cells/ml at a volume ratio of 1:1. From day 13 until the end of the experiment larvae were fed *C. muelleri*, and *I. galbana* at 100,000 cells/ml at a volume ratio of 3:1. Larvae entered their pediveliger stage and became competent to settle between 25-28 days. We deemed the larvae competent when they had a prominent eyespot and larval foot and measured 320-350 μm in diameter.

**Description of models used and post hoc results**

In experiment 1 the model included interactions effects between conspecific waterbourn and predator cues, between conspecific shells and waterbourn conspecific cues, and between predator cues and conspecific shells (see Table S2). In experiment 2 the model included interactions effects between conspecific waterbourn and predator cues, between conspecific waterbourn cues and biofilms, and between conspecific shells and biofilms (see Table S2). In the combined data, the model included interactions effects between conspecific waterbourn and predator cues, and between conspecific shells and waterbourn conspecific cues (see Table S2).

Table S3. Description of all the models used for each experiment as well as every interaction term. Models are uniquely identified so that they can be referenced for the post-hoc results.

| Experiment |  |
| --- | --- |
| Pilot study | glmer(Settled ~ predator_cue + conspecific_cue + predator_cue:conspecific_cue + Larvae.age + (1\|Larvae.batch), data = data, family = binomial) |
| Experiment 1 | glmer(Settled ~ conspecific_cue + predator_cue + Shell + conspecific_cue:predator_cue + conspecific_cue:Shell + predator_cue:Shell + Larvae.age + (1\|Larvae.batch), data=data, family=binomial) |
| Experiment 2 | glmer(Settled ~ conspecific_cue + Shell + biofilm + conspecific_cue:predator_cue + conspecific_cue:biofilm + Shell:biofilm + Larvae.age + (1\|Batch), data=data_aug, family=binomial) |
| Combined data | glmer(Settled ~ conspecific_cue + predator_cue + Shell + conspecific_cue:predator_cue + conspecific_cue:Shell + Larvae.age + (1 \| Larvae.batch), data = data_combined, family = binomial) |

Table S4. Results of the pairwise post hoc performed on every model.

| **Experiment** | **Contrasts** | | | | | **odds.ratio** | **SE** | **df** | **null** | **z.ratio** | **p.value** |
| --- | --- | --- | --- | --- | --- | --- | --- | --- | --- | --- | --- |
| Pilot | Conspecific_absent | Predator_absent | / | Conspecific_present | Predator_absent | 0.254 | 0.073 | Inf | 1 | -4.769 | <.0001 |
|  | Conspecific_absent | Predator_absent | / | Conspecific_absent | Predator_present | 0.895 | 0.294 | Inf | 1 | -0.339 | 0.9866 |
|  | Conspecific_absent | Predator_absent | / | Conspecific_present | Predator_present | 0.419 | 0.125 | Inf | 1 | -2.921 | 0.0183 |
|  | Conspecific_present | Predator_absent | / | Conspecific_absent | Predator_present | 3.528 | 0.973 | Inf | 1 | 4.571 | <.0001 |
|  | Conspecific_present | Predator_absent | / | Conspecific_present | Predator_present | 1.650 | 0.394 | Inf | 1 | 2.100 | 0.1531 |
|  | Conspecific_absent | Predator_present | / | Conspecific_present | Predator_present | 0.468 | 0.134 | Inf | 1 | -2.649 | 0.0402 |
| Experiment 1 | conspecific_absent | predator_absent | / | conspecific_present | predator_absent | 0.2279 | 0.0724 | Inf | 1 | -4.653 | <.0001 |
|  | conspecific_absent | predator_absent | / | conspecific_absent | predator_present | 2.1689 | 1.2482 | Inf | 1 | 1.345 | 0.5340 |
|  | conspecific_absent | predator_absent | / | conspecific_present | predator_present | 0.1920 | 0.1308 | Inf | 1 | -2.423 | 0.0728 |
|  | conspecific_present | predator_absent | / | conspecific_absent | predator_present | 9.5170 | 5.2264 | Inf | 1 | 4.103 | 0.0002 |
|  | conspecific_present | predator_absent | / | conspecific_present | predator_present | 0.8426 | 0.6030 | Inf | 1 | -0.239 | 0.9952 |
|  | conspecific_absent | predator_present | / | conspecific_present | predator_present | 0.0885 | 0.0982 | Inf | 1 | -2.187 | 0.1268 |
| Experiment 1 | Sterilized shell | predator_absent | / | Untreated shell | predator_absent | 0.04219 | 0.01375 | Inf | 1 | -9.714 | <.0001 |
|  | Sterilized shell | predator_absent | / | Sterilized shell | predator_present | 4.02126 | 2.38773 | Inf | 1 | 2.344 | 0.0884 |
|  | Sterilized shell | predator_absent | / | Untreated shell | predator_present | 0.01917 | 0.01266 | Inf | 1 | -5.990 | <.0001 |
|  | Untreated shell | predator_absent | / | Sterilized shell | predator_present | 95.31582 | 54.1249 | Inf | 1 | 8.025 | <.0001 |
|  | Untreated shell | predator_absent | / | Untreated shell | predator_present | 0.45446 | 0.30478 | Inf | 1 | -1.176 | 0.6421 |
|  | Sterilized shell | predator_present | / | Untreated shell | predator_present | 0.00477 | 0.00517 | Inf | 1 | -4.930 | <.0001 |
| Experiment 1 | sterilized | conspecific_absent | / | Untreated shell | conspecific_absent | 0.01259 | 0.00730 | Inf | 1 | -7.540 | <.0001 |
|  | sterilized | conspecific_absent | / | Sterilized shell | conspecific_present | 0.12610 | 0.07482 | Inf | 1 | -3.490 | 0.0027 |
|  | sterilized | conspecific_absent | / | Untreated shell | conspecific_present | 0.00201 | 0.00218 | Inf | 1 | -5.738 | <.0001 |
|  | untreated | conspecific_absent | / | Sterilized shell | conspecific_present | 10.01535 | 2.73373 | Inf | 1 | 8.441 | <.0001 |
|  | untreated | conspecific_absent | / | Untreated shell | conspecific_present | 0.16001 | 0.11035 | Inf | 1 | -2.657 | 0.0394 |
|  | sterilized | conspecific_present | / | Untreated shell | conspecific_present | 0.01598 | 0.01111 | Inf | 1 | -5.947 | <.0001 |
| Experiment 2 | conspecific_absent | predator_absent | / | conspecific_present | predator_absent | 0.212 | 0.0332 | Inf | 1 | -9.922 | <.0001 |
|  | conspecific_absent | predator_absent | / | conspecific_absent | predator_present | 0.932 | 0.1409 | Inf | 1 | -0.469 | 0.9659 |
|  | conspecific_absent | predator_absent | / | conspecific_present | predator_present | 0.300 | 0.0451 | Inf | 1 | -7.998 | <.0001 |
|  | conspecific_present | predator_absent | / | conspecific_absent | predator_present | 4.386 | 0.6875 | Inf | 1 | 9.431 | <.0001 |
|  | conspecific_present | predator_absent | / | conspecific_present | predator_present | 1.410 | 0.2114 | Inf | 1 | 2.292 | 0.0998 |
|  | conspecific_absent | predator_present | / | conspecific_present | predator_present | 0.322 | 0.0487 | Inf | 1 | -7.493 | <.0001 |
| Experiment 2 | conspecific_absent | biofilm_absent | / | conspecific_present | biofilm_absent | 0.207 | 0.0332 | Inf | 1 | -9.819 | <.0001 |
|  | conspecific_absent | biofilm_absent | / | conspecific_absent | biofilm_present | 0.372 | 0.0588 | Inf | 1 | -6.258 | <.0001 |
|  | conspecific_absent | biofilm_absent | / | conspecific_present | biofilm_present | 0.123 | 0.0198 | Inf | 1 | -12.999 | <.0001 |
|  | conspecific_present | biofilm_absent | / | conspecific_absent | biofilm_present | 1.794 | 0.2656 | Inf | 1 | 3.949 | 0.0005 |
|  | conspecific_present | biofilm_absent | / | conspecific_present | biofilm_present | 1.794 | 0.2656 | Inf | 1 | -3.491 | 0.0027 |
|  | conspecific_absent | biofilm_present | / | conspecific_present | biofilm_present | 0.591 | 0.0891 | Inf | 1 | -7.281 | <.0001 |
| Experiment 2 | conspecific_absent | Sterilized shell | / | conspecific_present | Sterilized shell | 0.261 | 0.02900 | Inf | 1 | -12.092 | <.0001 |
|  | conspecific_absent | Sterilized shell | / | conspecific_absent | Untreated shell | 0.130 | 0.01468 | Inf | 1 | -18.073 | <.0001 |
|  | conspecific_absent | Sterilized shell | / | conspecific_present | Untreated shell | 0.034 | 0.00607 | Inf | 1 | -18.934 | <.0001 |
|  | conspecific_present | Sterilized shell | / | conspecific_absent | Untreated shell | 0.497 | 0.06713 | Inf | 1 | -5.174 | <.0001 |
|  | conspecific_present | Sterilized shell | / | conspecific_present | Untreated shell | 0.130 | 0.01468 | Inf | 1 | -18.073 | <.0001 |
|  | conspecific_absent | Untreated shell | / | conspecific_present | Untreated shell | 0.261 | 0.02900 | Inf | 1 | -12.092 | <.0001 |
| Experiment 2 | predator_absent | biofilm_absent | / | predator_present | biofilm_absent | 1.146 | 0.1220 | Inf | 1 | 1.281 | 0.5749 |
|  | predator_absent | biofilm_absent | / | predator_absent | biofilm_present | 0.469 | 0.0507 | Inf | 1 | -7.007 | <.0001 |
|  | predator_absent | biofilm_absent | / | predator_present | biofilm_present | 0.537 | 0.0797 | Inf | 1 | -4.188 | 0.0002 |
|  | predator_present | biofilm_absent | / | predator_absent | biofilm_present | 0.409 | 0.0634 | Inf | 1 | -5.763 | <.0001 |
|  | predator_present | biofilm_absent | / | predator_present | biofilm_present | 0.469 | 0.0507 | Inf | 1 | -7.007 | <.0001 |
|  | predator_absent | biofilm_present | / | predator_present | biofilm_present | 1.146 | 0.1220 | Inf | 1 | 1.281 | 0.5749 |
| Experiment 2 | predator_absent | Sterilized shell | / | predator_present | Sterilized shell | 1.146 | 0.1220 | Inf | 1 | 1.281 | 0.5749 |
|  | predator_absent | Sterilized shell | / | predator_absent | Untreated shell | 0.130 | 0.0147 | Inf | 1 | -18.073 | <.0001 |
|  | predator_absent | Sterilized shell | / | predator_present | Untreated shell | 0.149 | 0.0229 | Inf | 1 | -12.384 | <.0001 |
|  | predator_present | Sterilized shell | / | predator_absent | Untreated shell | 0.113 | 0.0178 | Inf | 1 | -13.902 | <.0001 |
|  | predator_present | Sterilized shell | / | predator_present | Untreated shell | 0.130 | 0.0147 | Inf | 1 | -18.073 | <.0001 |
|  | predator_absent | Untreated shell | / | predator_present | Untreated shell | 1.146 | 0.1220 | Inf | 1 | 1.281 | 0.5749 |
| Experiment 2 | sterilized | biofilm_absent | / | Untreated shell | biofilm_absent | 0.106 | 0.0174 | Inf | 1 | -13.719 | <.0001 |
|  | sterilized | biofilm_absent | / | Sterilized shell | biofilm_present | 0.383 | 0.0618 | Inf | 1 | -5.944 | <.0001 |
|  | sterilized | biofilm_absent | / | Untreated shell | biofilm_present | 0.061 | 0.0102 | Inf | 1 | -16.642 | <.0001 |
|  | untreated | biofilm_absent | / | Sterilized shell | biofilm_present | 3.607 | 0.5176 | Inf | 1 | 8.940 | <.0001 |
|  | untreated | biofilm_absent | / | Untreated shell | biofilm_present | 0.574 | 0.0852 | Inf | 1 | -3.739 | 0.0011 |
|  | sterilized | biofilm_present | / | Untreated shell | biofilm_present | 0.159 | 0.0244 | Inf | 1 | -11.990 | <.0001 |
| Combined data | Conspecific waterbourn_absent | predator_absent | / | Conspecific waterbourn_present | predator_absent | 0.219 | 0.0298 | Inf | 1 | -11.142 | <.0001 |
|  | Conspecific waterbourn_absent | predator_absent | / | Conspecific waterbourn_absent | predator_present | 0.938 | 0.1243 | Inf | 1 | -0.479 | 0.9637 |
|  | Conspecific waterbourn_absent | predator_absent | / | Conspecific waterbourn_present | predator_present | 0.338 | 0.0474 | Inf | 1 | -7.738 | <.0001 |
|  | Conspecific waterbourn_present | predator_absent | / | Conspecific waterbourn_absent | predator_present | 4.286 | 0.5838 | Inf | 1 | 10.686 | <.0001 |
|  | Conspecific waterbourn_present | predator_absent | / | Conspecific waterbourn_present | predator_present | 1.588 | 0.2113 | Inf | 1 | 3.178 | 0.0081 |
|  | Conspecific waterbourn_absent | predator_present | / | Conspecific waterbourn_present | predator_present | 0.360 | 0.0505 | Inf | 1 | -7.286 | <.0001 |
| Combined data | Sterilized shell | predator_absent | / | untreated | predator_absent | 0.0964 | 0.00991 | Inf | 1 | -22.765 | <.0001 |
|  | Sterilized shell | predator_absent | / | Sterilized shell | predator_present | 1.2039 | 0.11459 | Inf | 1 | 1.950 | 0.2073 |
|  | Sterilized shell | predator_absent | / | Untreated shell | predator_present | 0.1161 | 0.01638 | Inf | 1 | -15.262 | <.0001 |
|  | Untreated shell | predator_absent | / | Sterilized shell | predator_present | 12.4847 | 1.73542 | Inf | 1 | 18.161 | <.0001 |
|  | Untreated shell | predator_absent | / | Untreated shell | predator_present | 1.2039 | 0.11459 | Inf | 1 | 1.950 | 0.2073 |
|  | Sterilized shell | predator_present | / | Untreated shell | predator_present | 0.0964 | 0.00991 | Inf | 1 | -22.765 | <.0001 |
| Combined data | Sterilized shell | conspecific_absent | / | Untreated shell | conspecific_absent | 0.0964 | 0.00991 | Inf | 1 | -22.765 | <.0001 |
|  | Sterilized shell | conspecific_absent | / | Sterilized shell | conspecific_present | 0.2809 | 0.02814 | Inf | 1 | -12.676 | <.0001 |
|  | Sterilized shell | conspecific_absent | / | Untreated shell | conspecific_present | 0.0271 | 0.00447 | Inf | 1 | -21.873 | <.0001 |
|  | Untreated shell | conspecific_absent | / | Sterilized shell | conspecific_present | 2.9126 | 0.34413 | Inf | 1 | 9.048 | <.0001 |
|  | Untreated shell | conspecific_absent | / | Untreated shell | conspecific_present | 0.2809 | 0.02814 | Inf | 1 | -12.676 | <.0001 |
|  | Sterilized shell | conspecific_present | / | Untreated shell | conspecific_present | 0.0964 | 0.00991 | Inf | 1 | -22.765 | <.0001 |
